# DUX4 is a common driver of immune evasion and immunotherapy failure in metastatic cancers

**DOI:** 10.1101/2023.07.10.548412

**Authors:** Jose Mario Bello Pineda, Robert K. Bradley

## Abstract

Cancer immune evasion contributes to checkpoint immunotherapy failure in many patients with metastatic cancers. The embryonic transcription factor DUX4 was recently characterized as a suppressor of interferon-γ signaling and antigen presentation that is aberrantly expressed in a small subset of primary tumors. Here, we report that *DUX4* expression is a common feature of metastatic tumors, with ∼10-50% of advanced bladder, breast, kidney, prostate, and skin cancers expressing *DUX4*. *DUX4* expression is significantly associated with immune cell exclusion and decreased objective response to PD-L1 blockade in a large cohort of urothelial carcinoma patients. *DUX4* expression is a significant predictor of survival even after accounting for tumor mutational burden and other molecular and clinical features in this cohort, with *DUX4* expression associated with a median reduction in survival of over one year. Our data motivate future attempts to develop DUX4 as a biomarker and therapeutic target for checkpoint immunotherapy resistance.

## INTRODUCTION

Immune checkpoint inhibition (ICI) therapy utilizes immunomodulatory monoclonal antibodies to stimulate patient anti-tumor immune responses. Blockade of T cell co-inhibitory receptors, such as CTLA-4 and the PD-1/PD-L1 axis, has achieved major success in the treatment of diverse metastatic cancers compared to first-line chemotherapy (Doki et al., 2022; Hellmann et al., 2019; Klein et al., 2020; Larkin et al., 2019; Motzer et al., 2020; Stein et al., 2022). However, a majority of advanced cancer patients fail to respond to ICI due to de novo or acquired resistance, the mechanistic bases of which remain incompletely understood.

Diverse mechanisms modulate sensitivity and resistance to immune checkpoint inhibition (Kalbasi & Ribas, 2020). These mechanisms include defects in MHC class I-mediated antigen presentation due to loss of *B2M* or *HLA* (Grasso et al., 2018; Lee et al., 2020; McGranahan et al., 2016; Sade-Feldman et al., 2017; Sucker et al., 2014; Wolf et al., 2019), *PTEN* and *LSD1* inactivation, which sensitizes tumor cells to type I interferon signaling (S. Li et al., 2016; Peng et al., 2016; Sheng et al., 2018), T cell dysfunction (Jiang et al., 2018), presence of specific T cell populations in the tumor microenvironment (Gide et al., 2019), and active WNT–β-catenin signaling (Spranger et al., 2015). MAPK signaling in *BRAF*-mutated melanomas and CDK4/CDK6 activity have also been implicated in reduced ICI efficacy, and combination treatment with a MAPK/CDK inhibitor improves response to checkpoint blockade (Ascierto et al., 2019; Deng et al., 2018; Ebert et al., 2016; Goel et al., 2017; Jerby-Arnon et al., 2018; Ribas et al., 2019; Schaer et al., 2018; Sullivan et al., 2019).

Tumor cell-intrinsic interferon-gamma (IFN-γ) signaling is particularly important in anti-tumor immunity. This pathway induces expression of genes involved in MHC class I-mediated antigen processing and presentation, which include genes encoding the TAP1/TAP2 transporters, components of the immunoproteasome, HLA proteins, and B2M (Alspach et al., 2019). Thus, suppression of IFN-γ activity promotes tumor immune evasion and decreased CD8^+^ T cell activation. Indeed, decreased ICI efficacy was observed in patients with tumors harboring inactivating mutations in IFN-γ pathway genes such as *JAK1* and *JAK2* (Gao et al., 2016; Nguyen et al., 2021; Sucker et al., 2017; Zaretsky et al., 2016). Similarly, a recent study reported a splicing-augmenting mutation in *JAK3*, linked to decreased *JAK3* expression levels, as a potential mechanism of resistance in a patient with metastatic melanoma treated with anti-PD-1 and anti-CTLA-4 combination therapy (Newell et al., 2022).

Some cancers exhibit aberrant expression of embryonic DUX transcription factors. For instance, *DUXB* is expressed in diverse primary malignancies, most notably in testicular germ cell and breast carcinomas (Preussner et al., 2018). Recent work from our group and others showed that *DUX4* is expressed in a small subset of primary tumors, where it suppresses tumor cell antigen presentation and response to IFN-γ signaling (Chew et al., 2019; Spens et al., 2023). We additionally observed signals that *DUX4* expression was associated with reduced survival following response to anti-CTLA-4 or anti-PD-1 in melanoma; however, those analyses relied upon two small cohorts (n = 27 or 41 patients), limiting the statistical power of our conclusions.

In its native embryonic context, DUX4 initializes human zygotic genome activation. *DUX4* expression levels peak at the 4-cell stage of the cleavage embryo; *DUX4* is then immediately silenced via epigenetic repression of the D4Z4 repeat array that contains the *DUX4* gene (de Iaco et al., 2017; Hendrickson et al., 2017; Himeda & Jones, 2019; Sugie et al., 2020; Whiddon et al., 2017). Aside from select sites of immune privilege such as the testis, *DUX4* remains silenced in adult somatic tissues (Das & Chadwick, 2016; Snider et al., 2010).

Since *DUX4* expression in cancer cells suppresses MHC class I-mediated antigen presentation (Chew et al., 2019), we hypothesized that *DUX4* expression might be particularly common in the setting of metastatic disease (versus the primary cancers that we studied previously), where immune evasion is particularly important. We therefore analyzed several large cohorts of patients with different metastatic cancers to determine the frequency of *DUX4* expression in advanced disease. We additionally rigorously tested the potential importance of *DUX4* expression for patient response to ICI in a well-powered cohort.

## RESULTS

### *DUX4* is commonly expressed in diverse metastatic cancer types

To assess the prevalence of *DUX4*-expressing human malignancies, we performed a large-scale analysis of publicly available RNA-seq data across diverse cancer types (**Fig. 1A, Fig. S1A**). The majority of the cohorts in The Cancer Genome Atlas (TCGA) are most commonly comprised of primary samples and local metastases. We found that *DUX4* expression is a particularly common feature across advanced-stage cancers, with 10-50% of cancer samples (depending upon cancer type) displaying *DUX4* expression levels comparable to or greater than those observed in the early embryo, where expression of the highly stereotyped DUX4-induced gene expression program is observed (Chew et al., 2019; Hendrickson et al., 2017). A markedly higher proportion of advanced metastatic cancers express *DUX4*—and tend to have higher absolute *DUX4* expression levels—than do their TCGA cancer counterparts (**Fig. 1B-C**).

**Figure 1.**
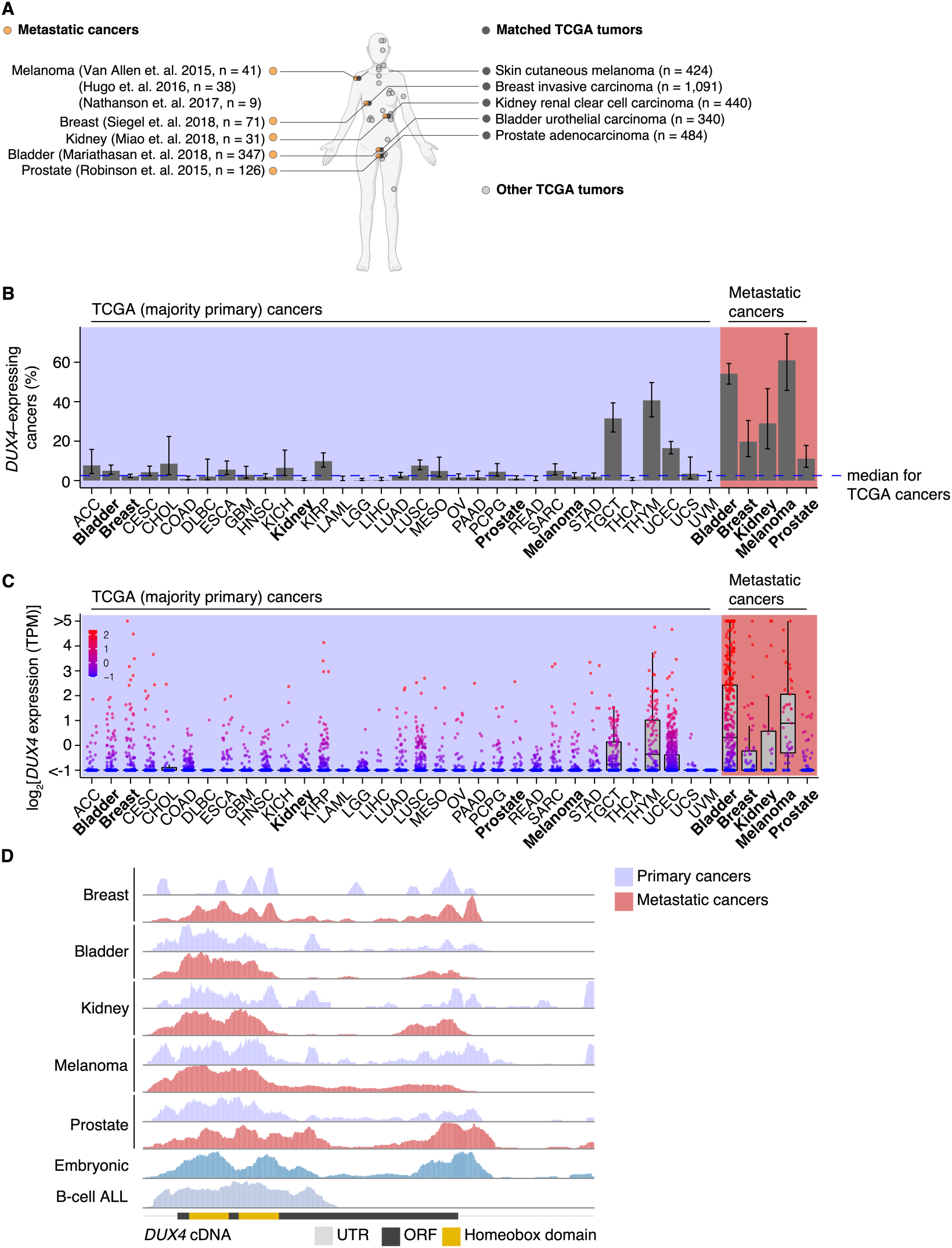
*DUX4* is frequently expressed in diverse metastatic cancers. (**A**) Matched The Cancer Genome Atlas (gray, TCGA) and advanced metastatic (orange) cancer datasets analyzed in our study. (**B**) The proportion of *DUX4*-expressing cancers in TCGA (purple shading) and metastatic (red shading) cancers. The blue line indicates the median over TCGA cancer cohorts. The 95% confidence intervals were estimated via a two-sided proportion test. (**C**) *DUX4* expression values (TPM, transcripts per million) in TCGA (purple shading) and advanced metastatic (red shading) cancer cohorts analyzed in our study. (**D**) Representative RNA-seq coverage plots from primary and metastatic cancers for reads mapping to the *DUX4* cDNA. Open reading frame (ORF, black rectangle); UTR (untranslated region, gray line); Homeobox domains (yellow rectangles).

We sought to determine if the *DUX4* transcripts in metastatic cancers express the entire coding sequence or only a portion thereof, as expressed *DUX4* truncations due to genomic rearrangements are frequent oncogenic drivers in particular cancer subtypes, most notably undifferentiated round cell sarcomas (CIC-DUX4 oncoprotein) (Antonescu et al., 2017; Choi et al., 2013; Graham et al., 2012; Italiano et al., 2012; Kawamura-Saito et al., 2006; Yoshida et al., 2016; Yoshimoto et al., 2009) and adolescent B-cell acute lymphoblastic leukemia (ALL) (Lilljebjörn et al., 2016; Liu et al., 2016; Qian et al., 2017; Yasuda et al., 2016). We aligned RNA-seq reads to the *DUX4* cDNA sequence and examined read coverage over the open reading frame. Resembling the cleavage stage embryo and *DUX4*-expressing primary cancers, *DUX4*-positive metastatic tumors transcribe the full-length coding region. In contrast, B-cell ALL exhibited the expected C-terminal truncation due to *DUX4* fusion with the *IGH* locus (**Fig. 1D**).

Since *DUX4* is typically silent in most healthy tissue contexts outside the cleavage-stage embryo (Das & Chadwick, 2016; Snider et al., 2010), we investigated if artifacts related to sequencing and sample processing could account for the observed high rates of *DUX4* expression in metastatic versus primary cancers. We were particularly interested in determining whether the method of RNA recovery influenced *DUX4* detection rate, as the analyzed metastatic cohorts frequently relied upon formalin-fixed samples rather than the frozen samples frequently used by TCGA. We took advantage of a cohort of patients with diverse metastatic tumor types for which patient-matched flash-frozen and formalin-fixed metastatic tumor samples were analyzed by RNA-seq (via poly(A)-selection and hybrid probe capture sequencing library preparations, respectively) (D. R. Robinson et al., 2017). Our re-analysis revealed that *DUX4* expression is readily detectable and quantifiable for both sample and library preparation methods. *DUX4* transcript levels in the majority of the sequenced samples were higher in poly(A)-selected sequencing than were the analogous measurements obtained from hybrid capture (**Fig. S1B-C**). These data demonstrate that the high rates of *DUX4* expression that we observed across metastatic cancer cohorts reflect true *DUX4* expression rather than technical biases introduced by studying formalin-fixed tissues and are consistent with expression of a polyadenylated *DUX4* transcript in both primary and metastatic cancers.

### *DUX4* expression is associated with immune cell exclusion

We next sought to assess the downstream consequences of *DUX4* expression in metastatic cancers. We focused on urothelial cancers for two reasons. First, urothelial cancers exhibited one of the highest frequencies of *DUX4* expression (54% of patients) in any of the five metastatic cancer cohorts that we analyzed, suggesting that DUX4 could be particularly important in that tumor type. Second, pretreatment samples from 347 patients enrolled in the IMvigor210 trial, a phase 2 trial of anti-PD-L1 (atezolizumab) therapy with advanced urothelial carcinoma, were subject to transcriptome profiling by RNA-seq as well as immunohistochemical analysis, enabling us to conduct comprehensive studies of the association between *DUX4* expression, the global transcriptome, and immunophenotypes in a well-powered cohort (Balar et al., 2017; Mariathasan et al., 2018; Rosenberg et al., 2016).

We examined associations between global gene expression profiles and *DUX4* expression in this advanced urothelial carcinoma cohort. We performed differential gene expression analyses on the individuals stratified according to tumor *DUX4* expression status. Gene Ontology (GO) network analyses on the upregulated genes in *DUX4*-positive cancers identified multiple clusters of development-associated terms, consistent with the known role of *DUX4* in early embryogenesis (**Fig. S2A;** de Iaco et al., 2017; Hendrickson et al., 2017; Sugie et al., 2020; Whiddon et al., 2017). By contrast, we found a single network associated with downregulated genes: GO terms corresponding to humoral or cell-mediated immunity (**Fig. 2A**). Using an IFN-γ gene signature predictive of response to blockade of the PD-1/PD-L1 axis, we found that *DUX4*-expressing cancers have statistically lower levels of IFN-γ activity (**Fig. S2B**; Ayers et al., 2017). Consistent with IFN-γ suppression, we observed extensive downregulation of genes involved in anti-tumor immunity such as those involved in MHC class I-dependent antigen presentation and T cell activation, checkpoint proteins, and chemokines involved in effector T cell recruitment. *DUX4*-expression was also correlated with suppression of genes critical for MHC class II-mediated antigen presentation, namely: MHC class II isotypes (*HLA–DP/DQ/DR)*, *HLA-DM* and *HLA-DO*, and the invariant chain (*CD74*) (Roche & Furuta, 2015). MHC class II gene expression is regulated by the transactivator CIITA via a conserved SXY-module present in the promoter regions of these genes. *CIITA* is induced by IFN-γ and is also conspicuously downregulated in *DUX4*-expressing tumors (**Fig. 2B**; (Glimcher & Kara, 1992; Masternak et al., 2000; Steimle et al., 1993, 1994). MHC class II-mediated antigen presentation can regulate T cell abundance in the tumor microenvironment and patient response to PD-1 blockade (Johnson et al., 2020). These analyses suggest that *DUX4* expression in the metastatic context induces an immunosuppressive gene expression program, concordant with its established function in inhibiting JAK-STAT signaling in primary cancers (Chew et al., 2019).

**Figure 2.**
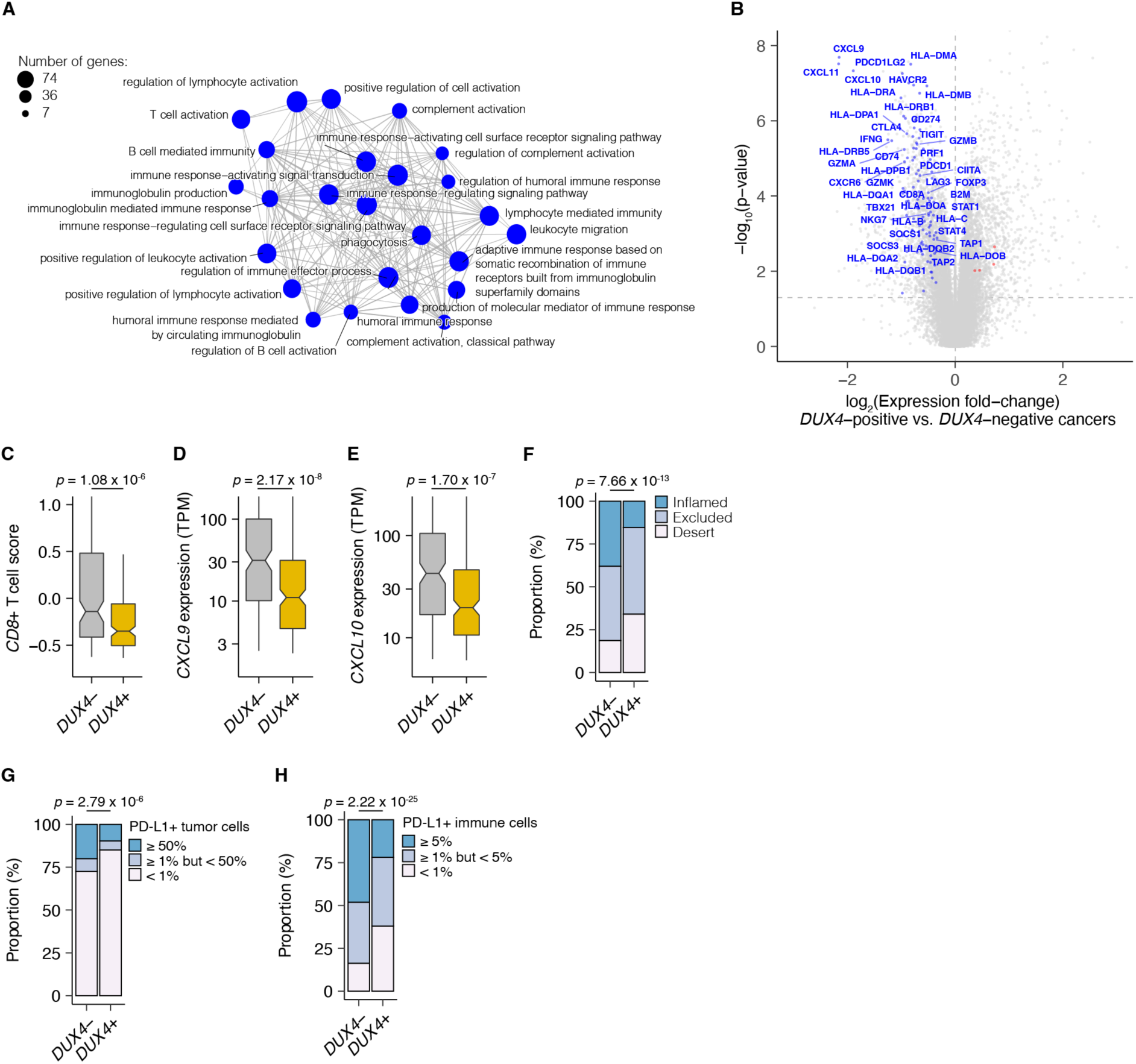
*DUX4* expression in advanced cancers is associated with signatures of host anti-tumor immunity inhibition. (**A**) Gene Ontology (GO) enrichment network analysis of *DUX4*-downregulated genes. Differentially expressed genes were identified from the comparison of advanced urothelial carcinoma tumors with high (> 1 TPM) vs. low (≤ 1 TPM) *DUX4* expression. The nodes and node sizes correspond to significantly enriched GO terms (Benjamini-Hochberg-adjusted *p*-value < 0.05) and the number of *DUX4*-downregulated genes in each, respectively. The edges connecting nodes indicate shared genes. (**B**) Downregulated (blue) and upregulated (red) anti-tumor immunity genes in tumors with *DUX4*-positive (> 1 TPM) vs. -negative (≤ 1 TPM) advanced urothelial carcinomas. (**C**) Effector CD8^+^ T cell score, defined as the mean of the z-score normalized gene expression values in the signature (Mariathasan et al., 2018) for *DUX4*+/- tumors. The *p*-value was estimated via a Mann-Whitney *U* test. (**D**) *CXCL9* expression for *DUX4*+/- tumors. The *p*-value was estimated via a Mann-Whitney *U* test. (**E**) As in (D), but illustrating *CXCL10* expression. (**F**) Proportion of immune phenotypes in *DUX4*+/- cancers. The phenotypes were based on the CD8^+^ T cell abundance and degree of tumor infiltration determined by anti-CD8 staining of tumor FFPE sections in the original study (Mariathasan et al., 2018). The *p*-value was estimated via a multinomial proportion test. (**G**) PD-L1-expression on tumor cells stratified by *DUX4* expression status measured by immunohistochemistry in the original study. The samples were categorized based on the percentage of PD-L1-positive tumor cells. The *p*-value was estimated via a multinomial proportion test. (**H**) As in (G), but PD-L1 staining on tumor-infiltrating immune cells (lymphocytes, macrophages, and dendritic cells) is represented.

We hypothesized that *DUX4* expression in these cancers will generate related transcriptomic signals consistent with CD8^+^ T cell exclusion from the tumor. We assessed this using an effector CD8^+^ T cell transcriptomic signature developed from initial studies of the IMvigor210 phase 2 trial (Balar et al., 2017; Rosenberg et al., 2016). *DUX4*-expressing cancers had lower measures of the gene signature, consistent with decreased CD8^+^ T cell infiltration into the tumor (**Fig. 2C**). We also investigated the possible effects of *DUX4* expression on the exclusion of other immune cell types using gene signatures developed from The Cancer Genome Atlas (Danaher et al., 2017). In these analyses, we recapitulated the observation of lower CD8^+^ T cell signature associated with *DUX4* positivity (**Fig. S2C)**. In addition, we observed patterns consistent with widespread immune cell exclusion from the tumor microenvironment (**Fig. S2D**). Defects in chemokine signaling could partially account for the observed *DUX4*-associated decrease in immune gene signature measurements. To test this hypothesis, we examined expression of chemokines involved in immune cell recruitment. In *DUX4*-expressing cancers, we observed lower mRNA levels of *CXCL9* and *CXCL10*, chemokines which recruit T cells to the tumor site (**Fig. 2D-E**; Nagarsheth et al., 2017). Utilizing a chemokine signature associated with host immune response to solid tumors, we observed that *DUX4* expression was correlated with broad reduction in the expression of chemokine signaling genes, beyond T cell-associated signals (**Fig. S2E**; Coppola et al., 2011).

We directly assessed the correlation of *DUX4* expression to immune cell exclusion by examining CD8^+^ T cell abundance in the tumor microenvironment, measured by immunohistochemistry (IHC) on formalin fixed paraffin embedded (FFPE) patient tumor sections. We verified that *DUX4* expression in the advanced urothelial carcinoma tumors was associated with an immune exclusion phenotype: a higher proportion of *DUX4*+ tumors exhibit either an immune-excluded or immune-desert phenotype compared to malignancies where *DUX4* is silent (**Fig. 2F**, **Fig. S2F**). We similarly examined the correlation of *DUX4* expression status with PD-L1 levels in the tumor and immune compartments quantified via IHC. We determined that *DUX4* expression was associated with a significant decrease in PD-L1 levels on both tumor and host immune cells, consistent with *DUX4*-induced suppression of IFN-γ signaling (**Fig. 2G**, **Fig. S2G, Fig. 2H**, **Fig. S2H**). PD-L1 expression on immune cells such as dendritic cells and macrophages modulate anti-tumor immune suppression and response to ICI in *in vivo* mouse models (Lau et al., 2017; Lin et al., 2018; Noguchi et al., 2017). Importantly, PD-L1 levels on immune cells are correlated with response to ICI in clinical trials (Powles et al., 2014; Rosenberg et al., 2016).

### DUX4 expression is correlated with poor response to immune checkpoint inhibition in advanced urothelial carcinoma

Given the correlation between cancer *DUX4* expression and signatures of anti-tumor immune response suppression, we sought to understand if *DUX4* expression in patient tumors was associated with accompanying changes to overall survival during PD-L1 inhibition. *DUX4* expression was associated with a significant decrease in objective response rates, assessed using the Response Evaluation Criteria in Solid Tumors (RECIST) (**Fig. 3A**). As expected, higher tumor mutational burden (TMB) was linked to improved survival outcomes in this cohort (**Fig. 3B**). Interestingly, we found that *DUX4* expression was correlated with statistically lower survival rates in this cohort after crudely adjusting for the effects of TMB by removing the bottom quartile of patients, those with the lowest number of missense mutations in their tumors (**Fig. 3C, Fig. S3A**). Those results motivated us to more carefully control for the effects of other patient covariates in order to better clarify the effects of *DUX4* expression on survival.

**Figure 3.**
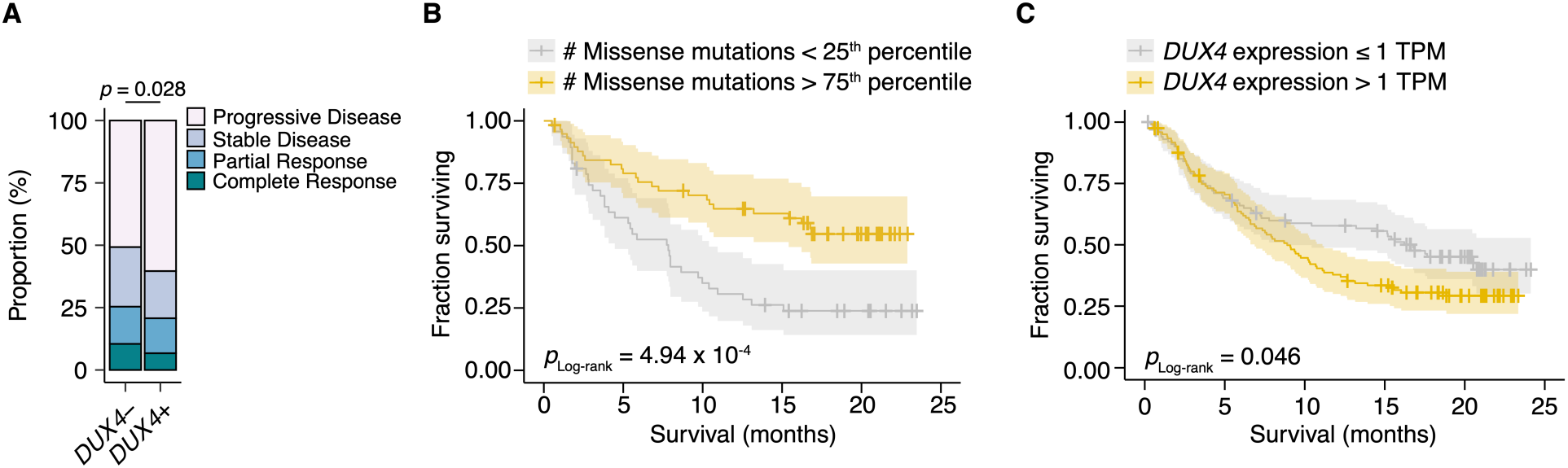
*DUX4*-positivity is associated with decreased response to immune checkpoint inhibition. (**A**) The proportion of clinical response classifications (RECIST, Response Evaluation Criteria in Solid Tumors) in *DUX4*-positive (*DUX4*+, > 1 TPM) or -negative (*DUX4*-, ≤ 1 TPM) advanced urothelial carcinoma patients. RECIST categories were assigned in the original study (Mariathasan et al., 2018). The *p*-value was estimated via a multinomial proportion test. (**B**) Kaplan-Meier (KM) estimates of overall survival for the patients in (A) stratified by tumor mutational burden (TMB, number of missense mutations). The estimated survival functions (solid lines), censored events (crosses), and 95% confidence intervals (transparent ribbons) for the patients in the top and bottom TMB quartiles are plotted. The *p*-value was estimated via a log-rank test. (**C**) As in (B), but patients are stratified by *DUX4* expression. To control for possible confounding by TMB, the quartile of patients with the lowest TMB was excluded.

### Risk assignments are improved with *DUX4* expression

We next determined whether *DUX4* expression was a significant predictor of survival for ICI-treated patients after controlling for TMB and other potentially relevant variables in a statistically rigorous manner. We used Cox Proportional Hazards (PH) regression to quantify the effects of multiple clinical, demographic, and molecular features on risk of death during ICI. For these and subsequent analyses, we elected to define *DUX4*-negative samples as those with *DUX4* expression levels < 0.25 TPM. This scheme excludes 126 patients but presumably produces more reliable categorizations, avoiding potential misclassifications due to loss of sensitivity of bulk RNA-Seq at very low expression levels (Mortazavi et al., 2008). In the context of multivariate Cox PH regression, which controls for the confounding effects of all other covariates simultaneously, we observed that TMB was positively associated with survival [hazard ratio (HR) = 0.14], as expected. Conversely, *DUX4* expression, Eastern Cooperative Oncology Group Performance Status (ECOG PS) > 0, and previous administration of platinum chemotherapy were correlated with increased risk (or shorter survival), while other features that have previously been reported as associated with reduced survival [e.g., *TGFB1* expression (Mariathasan et al., 2018)] did not remain significant after controlling for TMB and other variables. In particular, *DUX4* positivity was associated with dramatically worse survival, with a 3.2-fold increase in risk of death at any point in time compared to *DUX4*-negative status (**Fig. 4A**, **Table 1, Table S1**).

**Figure 4.**
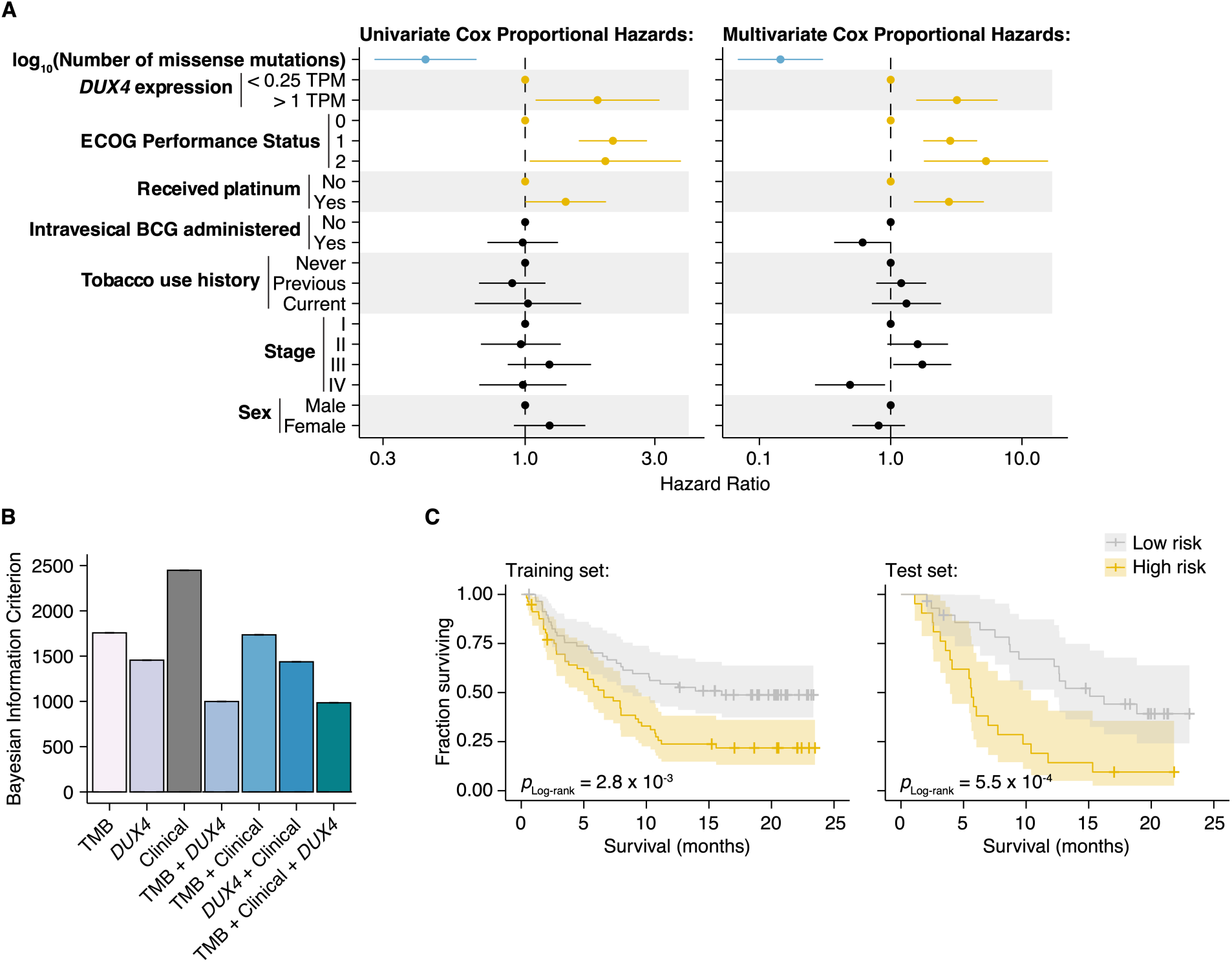
DUX4 expression status affects clinical response after controlling for other genetic and clinical variables. (**A**) Hazard ratios (HR) and 95% confidence intervals for the variables included in univariate (left) or multivariate (right) Cox Proportional Hazards (PH) regression. For categorical variables, the reference groups are indicated by points at HR = 1. Statistically significant predictors that are associated with increased (orange) or decreased (blue) risk in both the univariate and multivariate contexts are highlighted. ECOG (Eastern Cooperative Oncology Group); BCG (Bacillus Calmette-Guerin). (**B**) Bayesian information criterion (BIC) measurements for goodness of fit for the full (TMB, Clinical, DUX4 expression) vs. reduced Cox PH models, where lower values indicate better fit. The bootstrapped BIC mean and the 95% confidence interval are illustrated. Clinical (ECOG Performance Status and Platinum treatment history). (**C**) Kaplan-Meier (KM) estimates of overall survival, 95% confidence interval (transparent ribbon), and censored events (crosses) for low-risk (solid gray line) and high-risk (solid orange line) patients in the training (left) and test (right) sets. Risk group assignments were based on risk scores estimated by the full Cox PH model. *p*-values were estimated via a log-rank test.

**Table 1.** Cox Proportional Hazards Regression for Overall Survival.

| Characteristic | Univariate |  |  |  | Multivariate<sup>a,b</sup> |  |  |  |
| --- | --- | --- | --- | --- | --- | --- | --- | --- |
|  | HR | 95% CI | p-value | q-value<sup>c</sup> | HR | 95% CI | p-value | q-value<sup>c</sup> |
| Tumor mutational burden<sup>d</sup> | 0.43 | 0.28 - 0.66 | 1.10 x 10<sup>-4</sup> | 6.62 x 10<sup>-4</sup> | 0.14 | 0.07 - 0.30 | 2.88 x 10<sup>-7</sup> | 3.46 x 10<sup>-6</sup> |
| DUX4 expression: |  |  |  |  |  |  |  |  |
| < 0.25 TPM (reference) | — | — | — | — | — | — | — | — |
| > 1 TPM | 1.85 | 1.10 - 3.11 | 0.021 | 0.084 | 3.19 | 1.58 - 6.46 | 1.24 x 10<sup>-3</sup> | 3.71 x 10<sup>-3</sup> |
| ECOG Performance Status<sup>e</sup>: |  |  |  |  |  |  |  |  |
| 0 (reference) | — | — | — | — | — | — | — | — |
| 1 | 2.10 | 1.58 - 2.79 | 2.95 x 10<sup>-7</sup> | 3.54 x 10<sup>-6</sup> | 2.84 | 1.79 - 4.52 | 1.00 x 10<sup>-5</sup> | 6.02 x 10<sup>-5</sup> |
| 2 | 1.97 | 1.04 - 3.73 | 0.036 | 0.11 | 5.32 | 1.81 - 15.66 | 2.41 x 10<sup>-3</sup> | 5.79 x 10<sup>-3</sup> |
| Received platinum: |  |  |  |  |  |  |  |  |
| No (reference) | — | — | — | — | — | — | — | — |
| Yes | 1.41 | 1.01 - 1.98 | 0.047 | 0.11 | 2.78 | 1.52 - 5.08 | 9.19 x 10<sup>-4</sup> | 3.67 x 10<sup>-3</sup> |
| Intravesical BCG administered: |  |  |  |  |  |  |  |  |
| No (reference) | — | — | — | — | — | — | — | — |
| Yes | 0.98 | 0.73 - 1.32 | 0.89 | 0.92 | 0.61 | 0.37 - 1.01 | 0.054 | 0.080 |
| Tobacco use history: |  |  |  |  |  |  |  |  |
| Never (reference) | — | — | — | — | — | — | — | — |
| Previous | 0.896 | 0.68 - 1.19 | 0.45 | 0.67 | 1.20 | 0.78 - 1.86 | 0.41 | 0.41 |
| Current | 1.02 | 0.65 - 1.61 | 0.92 | 0.92 | 1.32 | 0.72 - 2.42 | 0.37 | 0.41 |
| Stage: |  |  |  |  |  |  |  |  |
| I | — | — | — | — | — | — | — | — |
| II | 0.96 | 0.69 - 1.35 | 0.83 | 0.92 | 1.60 | 0.62 - 0.94 | 0.082 | 0.11 |
| III | 1.23 | 0.86 - 1.75 | 0.25 | 0.44 | 1.74 | 1.05 - 2.90 | 0.033 | 0.056 |
| IV | 0.98 | 0.68 - 1.42 | 0.92 | 0.92 | 0.49 | 0.27 - 0.91 | 0.022 | 0.046 |
| Sex: |  |  |  |  |  |  |  |  |
| Male (reference) | — | — | — | — | — | — | — | — |
| Female | 1.23 | 0.91 - 1.66 | 0.18 | 0.36 | 0.81 | 0.51 - 1.29 | 0.37 | 0.41 |

<sup>a</sup>Log-rank test: p-value = 1.40 x 10<sup>-8</sup>
<sup>b</sup>Akaike Information Criterion = 970.78; Bayesian Information Criterion = 1003.08; Harrell's Concordance Index = 0.70
<sup>c</sup>Benjamini-Hochberg FDR correction
<sup>d</sup>log<sub>10</sub>(number of missense mutations)
<sup>e</sup>Eastern Oncology Cooperative Group Performance Status

We next investigated if *DUX4* expression status carried added value as a predictor over routinely collected clinical and molecular information. We focused on the variables with significant hazard ratios under both the univariate and multivariate regression settings: *DUX4* expression status, TMB, ECOG PS, and history of platinum chemotherapy. We employed goodness of fit measurements, which compare the observed data to expectations from Cox PH models created using various combinations of the covariates. In these analyses, we observed a quantifiable improvement in data-model congruence with the addition of *DUX4* expression status (**Fig. 4B**, **Fig. S4A-B**). Additionally, we measured statistically significant differences in the likelihoods of the reduced models (without *DUX4* expression as a predictor) when compared to the full model (employs all covariates) (**Table 2**). Taken together, these analyses indicate that *DUX4* expression status is an informative predictor of risk under ICI treatment.

**Table 2.**
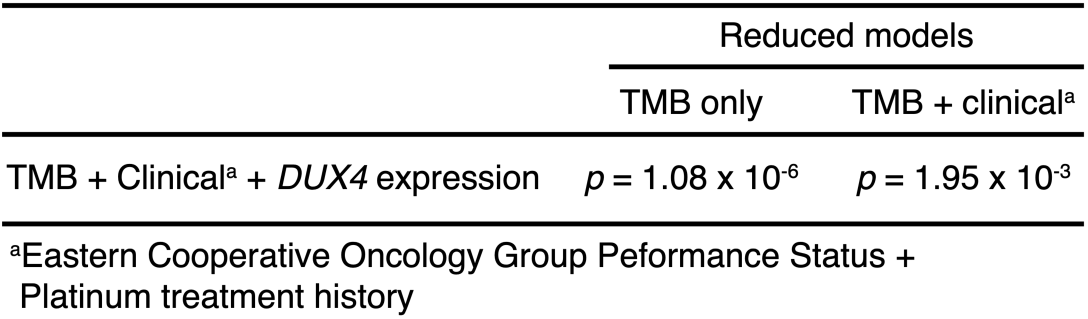
Likelihood ration rest.

We evaluated the utility of *DUX4* expression status for pre-treatment risk assignment in predicting patient response to ICI. We trained full and reduced Cox PH models on randomly sampled patients (training set, 70% of the cohort) and quantified their respective risk scores. A reference risk score per model was computed as the median score across the training set and was used to ascribe patients into low-vs. high-risk groups. Using these models, we quantified risk scores on the individuals excluded from model construction (test set, 30% of the patients), and similarly assigned patients into low-or high-risk groups based on the training set reference score. By empirically quantifying survival of the two risk groups using KM (Kaplan-Meier) estimation, we found that the full model stratifies patients in an informative manner, appropriately discriminating patients with longer vs. shorter survival times (**Fig. 4C**, **Fig. S4C-D**). Further, the addition of *DUX4* expression status improves model performance as illustrated by the time-dependent Brier score, a measure of survival prediction accuracy at specific timepoints **(Fig. S4E**).

### *DUX4* expression impedes response to ICI after controlling for other clinical characteristics

We used a Random Survival Forest (RSF) model to quantify the effect of *DUX4* expression on survival in ICI-treated advanced urothelial carcinoma patients (Ishwaran et al., 2008). The Random Survival Forest (RSF) is a machine learning ensemble, an extension of the Random Forest algorithm for right-censored data (Breiman, 2001). It can provide accurate estimates of risk and survival probability at definite times by aggregating predictions from a multitude of base learners (survival trees) (Ishwaran et al., 2008). RSFs have been successfully used to study time-to-event problems in medicine, including measurement of variable importance (Dietrich et al., 2016; Hsich et al., 2019; Ishwaran et al., 2009; O’Brien et al., 2021; Semeraro et al., 2011). We utilized the RSF model to address potential limitations of our Cox PH analyses. First, the RSF model is fully non-parametric and as such does not operate under the Cox PH assumptions: a constant relative hazard between strata over time (proportional hazards), a linear relationship between the predictors and the log hazard, and the unspecified baseline hazard function. Second, the RSF model can compute estimates of absolute risk and survival probability over time independent of a reference, unlike relative risk models such as Cox PH (Ishwaran et al., 2008).

We used all available molecular, clinical, and demographic covariates to grow an RSF. We randomly selected 70% of the patients to grow the forest, with the resulting model having an Out-of-Bag (OOB) error of 38.4%. The OOB error stabilizes with increasing number of trees and converges to the leave-one-out cross-validation error estimate. Thus, OOB error is characterized as an unbiased estimate of the model’s true prediction error (Breiman, 2001; Hastie et al., 2009). In some instances, the OOB error provides overestimates and some reports have recommended treating it as an upper bound (Bylander, 2002; Janitza & Hornung, 2018; Mitchell, 2011). Thus, we measured the RSF model’s test error using a holdout set (the remaining 30% of the cohort) excluded from training. The RSF model recorded a test error of 32.6% illustrating an appropriate fit (**Fig. S5A**). Our error measurements are comparable to Ishwaran, et al. (2008), suggesting our model can be used for inference purposes. Further, the time-dependent Brier score of the RSF model on the training and test sets confirms informative survival prediction (**Fig. S5B**).

The RSF model predicted worse survival outcomes in patients with *DUX4*-expressing cancers compared to their *DUX4*-silent counterparts. These predictions were mirrored in the test dataset, illustrating robustness of the model (**Fig. 5A**). Using time-dependent Receiver Operating Characteristic (ROC) curve analyses, we identified the time range for which the RSF predictive performance is statistically divergent from random guessing: approximately 6 to 20 months (**Fig. S5C**). In this window we observed significant survival differences between patients with *DUX4*+ and *DUX4*-tumors. We highlighted the model’s performance at predicting 1-year and 1.5-year survival, typical timepoints of clinical interest. For these times, the RSF appropriately discriminates patient death and survival (**Fig. S5D**). Examining the absolute effects of *DUX4* expression on survival, the RSF model predicted an approximately 20% decrease in both 1-year and 1.5-year survival probabilities in patients with *DUX4*-expressing cancers (**Fig. 5B**). Importantly, RSF survival predictions conform closely with the empirical survival estimates obtained via the Kaplan-Meier model (**Fig. 5C**).

**Figure 5.**
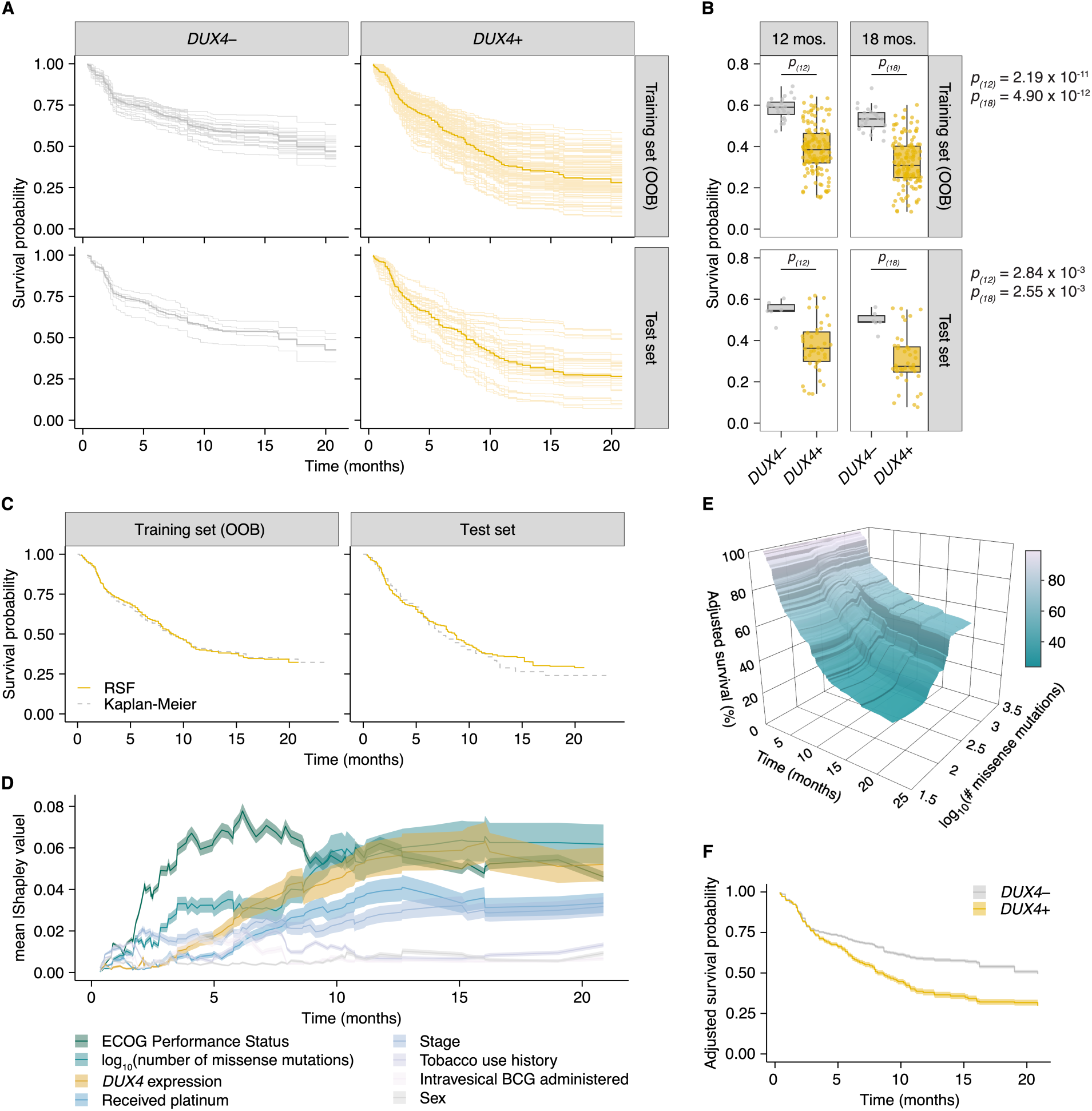
*DUX4* expression is associated with decreased overall survival in the context of immune checkpoint inhibition. (**A**) Random Survival Forest (RSF) predicted overall survival for patients with either *DUX4*-positive or -negative tumors in the training and test sets. Out-of-bag (OOB) survival predictions are shown for the patients in the training set. Survival predictions for individual patients (thin lines) and the median survival function across the cohort (thick line) are represented. *DUX4*-(< 0.25 TPM); *DUX4*+ (> 1 TPM). (**B**) Training (OOB) and test set survival probability predictions for patients with *DUX4*+/- tumors at 12 and 18 months. The *p*-values were estimated using a two-sided Mann-Whitney *U* test. (**C**) Survival probability for patients in the training and test sets. Survival functions corresponding to the median RSF prediction (solid orange line) and the Kaplan-Meier estimate (dashed gray) are displayed. (**D**) Feature importance for variables used in the RSF model. The average absolute estimated Shapley values (solid lines) are shown, associated with predicting survival probability at particular times. The 95% confidence interval of the mean (transparent ribbon) is plotted. (**E**) Surface plot showing adjusted (marginal) survival probability, measured via partial dependence, as a function of tumor mutational burden (TMB, number of missense mutations) and time. Each point on the surface corresponds to the mean survival prediction (at the respective timepoint) after TMB is fixed to the respective value for all patients. (**F**) Partial plot showing adjusted survival probability as a function of *DUX4* expression status. The median survival probability (solid lines) and the 95% confidence interval (transparent ribbon) after *DUX4* expression status is fixed to the indicated value for all patients are plotted.

We sought to determine the importance of *DUX4* expression status relative to the other covariates in the RSF model. We measured feature importance using estimated Shapley values, which quantify the marginal contribution of each variable to the RSF prediction (Lundberg & Lee, 2017; Maksymiuk et al., 2020; Shapley, 1953; Štrumbelj & Kononenko, 2014). Specifically, Shapley values measure variable contributions to predictions at the level of each patient. Contributions to the overall performance of the RSF model can be assessed by examining the aggregated summary: the average of the absolute Shapley values for a predictor across the patient cohort. We estimated Shapley values associated with predicting ensemble mortality, the RSF risk estimate. In these analyses, ECOG PS had the largest contribution, followed by TMB and *DUX4* expression (**Fig. S5E**). We validated these feature rankings using two independent metrics. The first metric was permutation importance, which quantifies the change in prediction error associated with permutation of a variable’s data; important covariates will record large deviations from the original predictions (Breiman, 2001; Ishwaran, 2007). The second measure employed was minimal depth, a measure of the variable-node-to-root-node distance within the survival trees of the RSF; important variables tend to have smaller minimal depth values as they are typically used for earlier decision splits (Ishwaran et al., 2010, 2011). Feature contributions measured using permutation importance and minimal depth were consistent with the Shapley-based assignments, notably identifying *DUX4* expression as an important contributor to patient survival outcomes (**Fig. S5E**). We investigated time-dependent changes in variable importance by estimating Shapley values associated with predicting survival probability at distinct time points along the observation window. Interestingly, we observed the strong dependence on ECOG PS for predicting survival at early timepoints under this paradigm. The importance of *DUX4* expression for survival prediction is most prominent at later times (**Fig. 5D**). Altogether, we found that diverse variable importance measures converge on identifying *DUX4* as a major contributor to patient survival prediction.

We sought to quantify the effect of *DUX4* expression on survival predictions after controlling for the effects of the other covariates. With Shapley dependence plots, which allows visualization of the marginal effects of a variable on the predicted outcome, we measured the expected negative correlation between TMB and mortality (**Fig. S5F**; Lundberg et al., 2020). We performed a similar dependence analysis on *DUX4* expression and observed a clear separation of positive and negative Shapley values based on *DUX4*-positive and -negative status, respectively. These results signify an increase in predicted risk of death associated with *DUX4* expression (**Fig. S5G**). Shapley values are scaled, variable attributions which add up to the difference between the prediction for an individual and the average prediction across the entire cohort and thus require transformation to get absolute measures of risk. To quantify the effects of TMB and *DUX4* expression in the appropriate risk units (mortality, expected number of deaths), we utilized partial dependence as an alternative way to represent mortality predictions as a function of these variables, marginalized over the other predictors in the data (Friedman, 2001). Specifically, the average model predictions across the individuals in the cohort are calculated over the unique predictor values. The marginal effects of TMB and *DUX4* expression measured via partial and Shapley dependence analyses show strong concordance. Comparison of the patients with the lowest versus highest TMB revealed an approximately 67% difference in mortality. On the other hand, we measured an approximately 53% increase in mortality associated with *DUX4*-positivity (**Fig. S5H-I**). We then extended the partial dependence analyses to survival probability predictions over time. In this paradigm, we similarly observed that higher TMB was correlated with increased survival probability, more pronounced at later times (**Fig. 5E**). *DUX4* expression was correlated with poorer survival outcomes, with a 1-year and 1.5-year survival difference of 20.7% and 19.2% between patients with *DUX4*+ and *DUX4*-tumors, respectively. Strikingly, our analyses measure a difference of at least 12.5 months in median survival between the *DUX4*+ and *DUX4*-strata (**Fig. 5F**). Overall, our analyses demonstrate a significant and robust decrease in survival attributable to *DUX4* expression in advanced cancers.

## DISCUSSION

*DUX4* expression is a common feature of metastasis and may be an important driver of immune evasion. While the mechanism governing *DUX4* de-repression in cancer remains to be elucidated, we show that *DUX4* expression in the metastatic context is associated with reduced anti-tumor immunity, mirroring previous observations in primary cancers and cancer cell line models (Chew et al., 2019), and is correlated with decreased patient survival under ICI treatment.

The prognostic value of IFN-γ activity (Ayers et al., 2017; Grasso et al., 2020; Newell et al., 2022) and its non-redundancy relative to TMB in terms of influencing ICI response is widely appreciated (Cristescu et al., 2018; Newell et al., 2022; Rozeman et al., 2021). For example, patients with advanced melanoma that is nonresponsive to anti-CTLA-4 or anti-PD-1/PD-L1 therapy have higher frequencies of genetic alterations associated with IFN-γ signaling defects compared to responsive patients (Gao et al., 2016; Nguyen et al., 2021; Sucker et al., 2017). DUX4 has been implicated in modifying IFN-γ activity through direct binding and inhibition of STAT1 via its C-terminal domain (Chew et al., 2019; Spens et al., 2023). Our sequence analyses show that *DUX4* transcripts in the metastatic context contain the full-length coding region, suggestive of an intact capability as a STAT1 suppressor. *DUX4*’s ubiquitous expression across metastatic cancers and our controlled survival analyses emphasize *DUX4* as an underappreciated contributor to ICI resistance.

Our RSF model allowed us to interrogate changes in variable importance over time. For instance, the contribution of *DUX4* expression to survival prediction is most prominent at later time points, suggesting principal effects on long-term survival. Intriguingly, these analyses revealed the outsize influence of ECOG PS, a measure of patient functional status, on survival at early timepoints relative to other patient covariates. ECOG PS negatively impacts patient survival during ICI therapy, inferred from our multivariate Cox PH analysis and from findings of the IMvigor210 clinical trial: patients with ECOG PS = 2 (n = 24) had a median overall survival of 8.1 months, lower than the subgroup with ECOG PS < 2 (n = 35) whose median survival was not reached during the observation period (Balar et al., 2017). Other studies have similarly reported poorer outcomes associated with ICI treatment in patients with high ECOG PS (Chalker et al., 2022; Krishnan et al., 2022; Petrillo et al., 2020; Sehgal et al., 2021). Altogether, these results possibly indicate the existence of co-occurring conditions in patients with higher degrees of disability, predisposing them to adverse effects associated with ICI treatment— comorbidities whose effects presumably manifest shortly after therapy commencement. Our data underscore the utility of time-dependent approaches in identifying covariate-linked survival effects which may not be apparent in a summary computed over the entire time period.

Our results may have broad implications for ICI treatment. First, *DUX4* expression may promote patient resistance in a wide array of ICI modalities. Our previous work showed that *DUX4* expression is associated with resistance to anti-CTLA-4 and anti-PD-1 therapies (Chew et al., 2019). In our current study, we comprehensively demonstrate that *DUX4* modulates patient response to PD-L1 blockade. We also report that *DUX4* expression in metastasis is correlated with downregulation of *TIGIT* (Zhang et al., 2018) and other immune checkpoints whose interception are currently under clinical investigation: *HAVCR2/TIM3* (NCT02608268; Dixon et al., 2021) and *LAG3* (NCT02658981; Amaria et al., 2022; Tawbi et al., 2022). Second, the pervasive expression of *DUX4* in all the metastatic cohorts we examined exhibits its potential as a pan-cancer biomarker. We show that binary categorization of patients according to *DUX4* expression status was sufficient to stratify patients according to ICI response. Screening for *DUX4* tumor expression, with binarized results such as through IHC using anti-DUX4 antibodies, could have clinical utility.

Our data motivate the investigation into *DUX4’s* potential to prognosticate response to ICI. However, our current study is limited by the availability of sufficiently-sized ICI-treated cohorts with associated patient data on relevant characteristics such as demographics and risk factors. Additional randomized trial data from diverse metastatic cancer cohorts, with adequate genomic and clinical data, is imperative. As these become available in the future, extending the analyses we have outlined in this study will be important to appraise *DUX4*’s definitive clinical relevance, contextualized amongst response-modifying clinical variables, in the use of immunotherapy in the treatment of metastatic cancer.

## METHODS

### Genome annotations, gene expression, and Gene Ontology (GO) enrichment analyses

A genome annotation was created through merging of the UCSC knownGene (Meyer et al., 2013), Ensembl 71 (Flicek et al., 2013), and MISO v2.0 (Katz et al., 2010) annotations for the hg19/GRCh37 assembly. Further, this annotation was expanded by generating all possible combinations of annotated 5’ and 3’ splice sites within each gene. RNA-seq reads were mapped to the transcriptome using RSEM v1.2.4 (B. Li & Dewey, 2011) calling Bowtie v1.0.0 (Langmead et al., 2009), with the option “-v 2.” TopHat v.2.0.8b (Trapnell et al., 2009) was used to map the unaligned reads to the genome and to the database of splice junctions obtained from the annotation merging described previously. Gene expression estimates (TPM, transcripts per million) obtained were normalized using the trimmed mean of M values (TMM) method (M. D. Robinson & Oshlack, 2010). Endogenous expression of *DUX4* during early embryogenesis range from approximately 2 to 10 TPM (Chew et al., 2019; Hendrickson et al., 2017). We have therefore defined DUX4-positive samples as those with expression levels > 1 TPM. In the differential gene expression analyses for the DUX4-positive vs. -negative comparison, gene expression values per sample group were compared using a two-sided Mann-Whitney *U* test. Differentially expressed genes illustrated in Figure 2B were identified as those with an absolute log_2_(fold-change) ≥ log_2_(1.25) and a *p*-value < 0.05. GO enrichment analyses, using the clusterProfiler package (Wu et al., 2021; Yu et al., 2012), were performed on *DUX4*-upregulated or -downregulated genes [absolute log_2_(fold-change) ≥ log_2_(1.5) and a *p*-value < 0.05] compared against the set of coding genes. Significant GO terms were defined as “Biological Process” terms with a Benjamini-Hochberg FDR-adjusted *p*-value < 0.05. The top 25 significant GO terms were illustrated (Fig. 2A and Fig. S2A). To investigate *DUX4* RNA-seq coverage patterns, a fasta file containing the *DUX4* cDNA sequence was assembled, indexed using samtools (H. Li et al., 2009), and used as a reference for read pseudoalignment by kallisto v.0.46.1 (Bray et al., 2016). The following kallisto parameters were used: kmer size of 31, estimated fragment length of 200, and estimated fragment length standard deviation of 80. Usage of the single-end option (“-- single”) and bias correction (“--bias”) were also specified. *DUX4* read coverage was visualized using the Integrative Genomics Viewer (IGV, Thorvaldsdóttir et al., 2013).

### Gene signature analyses

For a given gene set, z-score normalization of the expression values per gene was performed across the patient cohort. The signature score was defined as the mean of the normalized values across the genes of the set.

### Survival analyses, goodness of fit measures, and risk modeling

Kaplan-Meier (KM) estimation, *p*-value estimates from the log-rank test, and Cox Proportional Hazards (PH) regression in the univariate and multivariate contexts were performed using the survival package (T. Therneau, 2022; T. M. Therneau & Grambsch, 2000). Goodness of fit evaluations of the Cox PH models were done by measuring the Akaike Information Criterion (AIC) and Bayesian Information Criterion (BIC). AIC and BIC metrics balance model complexity with maximized likelihood, penalizing feature number increases without a concomitant improvement in performance. The likelihood ratio test was also used to compare goodness of fit of full (all variables) vs. reduced (subset of variables) Cox PH models. Specifically, the null hypothesis that the simple model provides as good as a fit as the more complex model was evaluated. The AIC, BIC, and likelihood ratio test *p*-values were computed using R’s stats package (R Core Team, 2022). For the Cox PH risk modeling, the patients were randomly assigned into training (70%) and test (30%) datasets. The createDataPartition() function from the caret package (Kuhn, 2022) was used to preserve the *DUX4* status class distribution after splitting. Full and reduced Cox PH models were created using the training data and the risk scores for each respective model were calculated using caret’s predict.coxph() function. For a given patient, the calculated risk score is equal to the hazard ratio relative to a “reference patient” (an individual whose covariate values are set to the respective means, from the training set). Specifically, the risk score is the quotient of the patient’s and the reference’s exponentiated linear predictors (the sum of the covariates in the model, weighted by the model’s regression coefficients). A “reference risk score” for each model was defined as the median risk score in the training data. Patients were assigned into low- or high-risk groups if their risk scores were lower or higher than the reference, respectively. The trained models were used to calculate risk scores and assign risk labels (based on the training set risk score reference) in the test set. The survival difference between low- and high-risk patients were empirically assessed via KM estimation and the log-rank test. Visualizations were created using the ggplot2 (Wickham, 2016), dplyr (Wickham et al., 2022), and survminer (Kassambara et al., 2021) packages.

### Random Survival Forest, feature importance, and partial dependence

We implemented a Random Survival Forest (RSF) model, an ensemble of multiple base learners (survival trees), using the randomForestSRC package (Ishwaran et al., 2008). The RSF algorithm is an extension of the Random Forest Algorithm (Breiman, 2001) for usage with right-censored data. Here, B bootstrap datasets are created from the original data, used to grow B concomitant survival trees (usually constrained by membership size in the terminal nodes) constructed using a randomly selected subset of the variables. Terminal node statistics are obtained for each tree: the survival function (via the Kaplan-Meier estimator), the cumulative hazard function (CHF, via the Nelson-Aalen estimator), and mortality (expected number of deaths; sum of the CHF over time). The RSF prediction is the average across the forest. Of note, each bootstrap dataset excludes 36.8% of the original data on average, the out-of-bag (OOB) samples. Thus, predictions for a particular sample can be made using the subset of the trees for which it was excluded from training (OOB predictions). Similarly, the associated OOB error for the RSF model can calculated, representing an unbiased estimate of the test error. We randomly assigned patients into training (70%) and test (30%) datasets. Since the *DUX4*-positive status was a minority class, we utilized the createDataPartition() function from the caret package (Kuhn, 2022) to preserve the class distribution within the splits. To determine optimal hyperparameters, we evaluated 5,616 RSF models representing different combinations of ntree (number of trees), nodesize (minimum terminal node size), mtry (number of randomly selected splitting variables), na.action (handling of missing data), splitrule (splitting rule), and samptype (type of bootstrap). We selected the model with hyperparameters which minimized both the OOB training and the test errors (defined as 1 – concordance index), namely: ntree = 1500, nodesize = 15, mtry = 3, na.action = "na.impute", splitrule = "bs.gradient", and samptype = "swr." We specified the use of an nsplit (number of random splits) value of 0 to indicate evaluation of all possible split points and usage of the optimum. For test set predictions, patients with missing data were omitted (na.action = "na.omit").

Feature importance in the final RSF model was evaluated using 3 metrics. First, permutation importance was measured using randomForestSRC’s subsample() function. RSF permutation importance utilizes OOB values: a variable’s OOB data is permuted and the change in the new vs. original OOB prediction error is quantified. The RSF permutation importance values were standardized by dividing by the variance and multiplying by 100, and the variance and confidence regions were obtained via the delete-*d* jackknife estimator (Ishwaran & Lu, 2019). Second, the tree-based feature importance metric minimal depth was calculated using randomForestSRC’s var.select() function. The minimal depth threshold (mean minimal depth) is the tree-averaged threshold (conservative = “medium”). Last, Shapley values were estimated using the fastshap package (Greenwell, 2021), using 1000 Monte Carlo repetitions. For each prediction, the sum of the estimated Shapley values was corrected (adjust = TRUE) to satisfy the efficiency (or local accuracy) property: for an individual *i*, the sum of *i*’s feature contributions equal the difference between the prediction for *i* and the average prediction across the entire cohort. For the overall measure of importance, the Shapley values were estimated from the mortality predictions from the RSF model (Fig S5E). Mortality is defined as the number of expected deaths over the observation window. That is, if all patients in the cohort shared the same covariate values as patient *i* who has mortality *m_i_*, then an average of *m* deaths is expected (Ishwaran et al., 2008). For the time-dependent implementation, we estimated Shapley values associated with the per timepoint RSF survival probability predictions along the observation window (Fig 5C).

The relationships of *DUX4* expression and TMB to mortality or survival probability (marginal contributions) were assessed via Shapley dependence plots and partial dependence plots. Partial dependence values were obtained using randomForestSRC’s partial() function and OOB predictions for mortality and survival probability were used as input. Visualizations were created in the R programming environment using the dplyr (Wickham et al., 2022), ggplot2 (Wickham, 2016), pammtools (Bender & Scheipl, 2018), and plotly (Sievert, 2020) packages.

### Measuring survival model predictive accuracy

The time-dependent Receiver Operating Characteristic (ROC) curve analyses were done to evaluate the RSF model’s accuracy in differentiating patients who die before a particular time *t*, from those who survive past *t* (Heagerty & Zheng, 2005). Specifically, for each timepoint, the cumulative/dynamic Area Under the ROC curve (AUC^C/D^) was calculated by computing the sensitivity (true positive rate) and specificity (1 – false positive rate) associated with using RSF-predicted mortality as the prognostic marker. The time-dependent AUC^C/D^ and 95% confidence interval per time point were estimated using the timeROC package, which adds the inverse-probability-of-censoring weights (IPCW) to the sensitivity calculation to correct for selection bias due to right-censoring (Blanche et al., 2013). The out-of-bag (training) or the test mortality predictions were used as input. The time-dependent Brier score and the Continuous Ranked Probability Score (CRPS, integrated Brier score divided by time) for the Cox PH models were computed using the pec package (Mogensen et al., 2012). The time-dependent Brier score and the CRPS for the RSF model was calculated using the randomForestSRC package (Ishwaran et al., 2008). The Kaplan-Meier estimator for the censoring times was used to estimate the IPCW (cens.model = “marginal”). Harrell’s concordance index for the Cox PH and RSF models were calculated using the survival (T. Therneau, 2022; T. M. Therneau & Grambsch, 2000) and randomForestSRC packages, respectively. Visualizations were created in the R programming environment using the dplyr (Wickham et al., 2022) and ggplot2 (Wickham, 2016) packages.

## ACKNOWLEDGEMENTS

R.K.B. was supported in part by the NIH/NCI (R01 CA251138), NIH/NHLBI (R01 HL128239, R01 HL151651), and the Blood Cancer Discoveries Grant program through the Leukemia & Lymphoma Society, Mark Foundation for Cancer Research, and Paul G. Allen Frontiers Group (8023-20). R.K.B is a Scholar of The Leukemia & Lymphoma Society (1344-18) and holds the McIlwain Family Endowed Chair in Data Science. The results shown here are in part based upon data generated by the TCGA Research Network: https://cancergenome.nih.gov/.

## AUTHOR CONTRIBUTIONS

J.M.B.P. and R.K.B. designed the study, analyzed the data, and wrote the paper.

---

**Figure S1.**
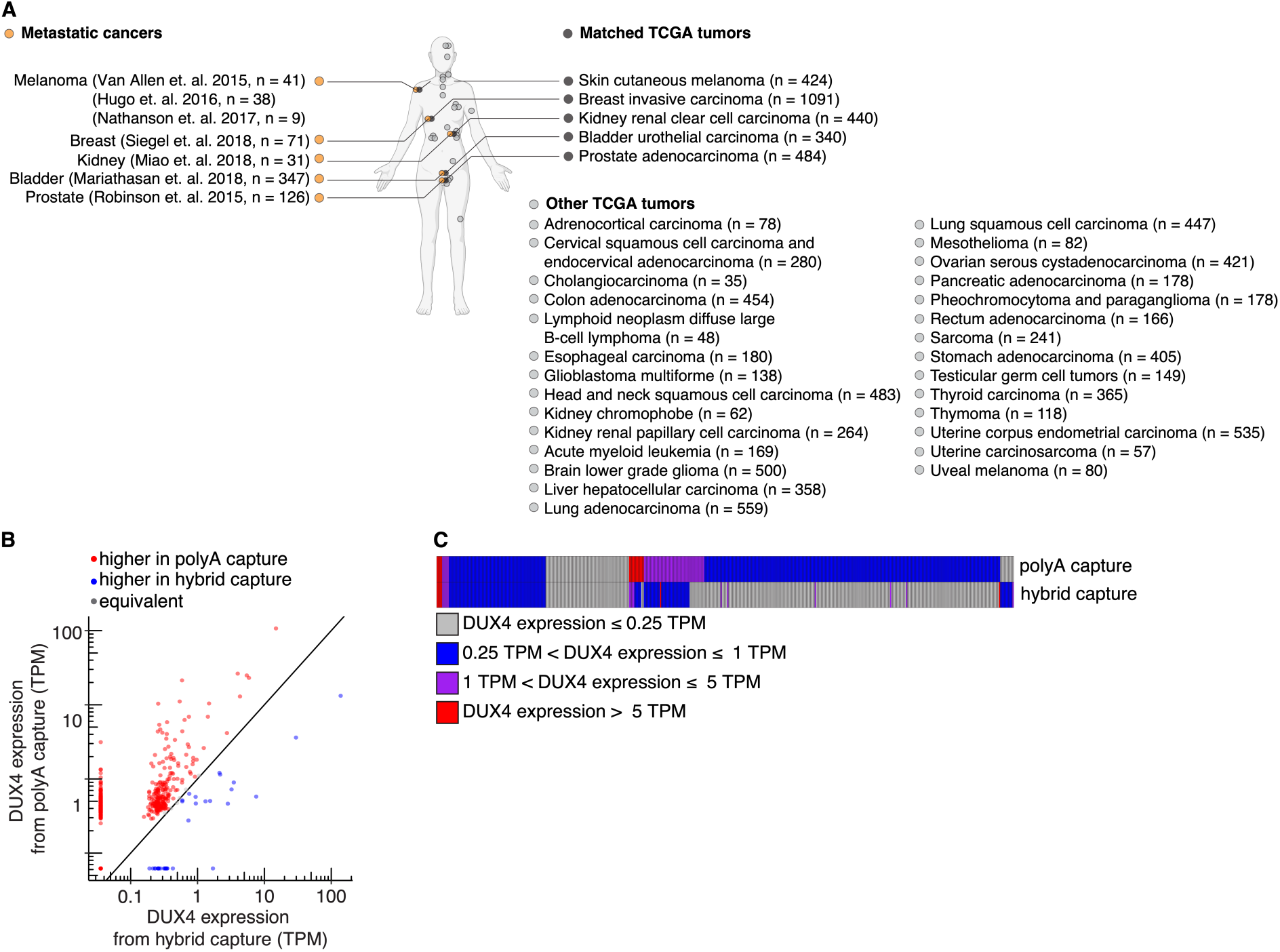
The *DUX4* transcript is likely polyadenylated. (**A**) As in (Figure 1A), but The Cancer Genome Atlas (TCGA) cancer cohorts without matched advanced metastatic counterparts analyzed in our study are shown. (**B**) A comparison of *DUX4* expression values (TPM, transcripts per million) measured from sequencing libraries prepared via poly(A) capture or hybrid capture. (**C**) As in (B), but a heatmap where patient samples (columns) were stratified according to the indicated categories of *DUX4* expression.

**Figure S2.**
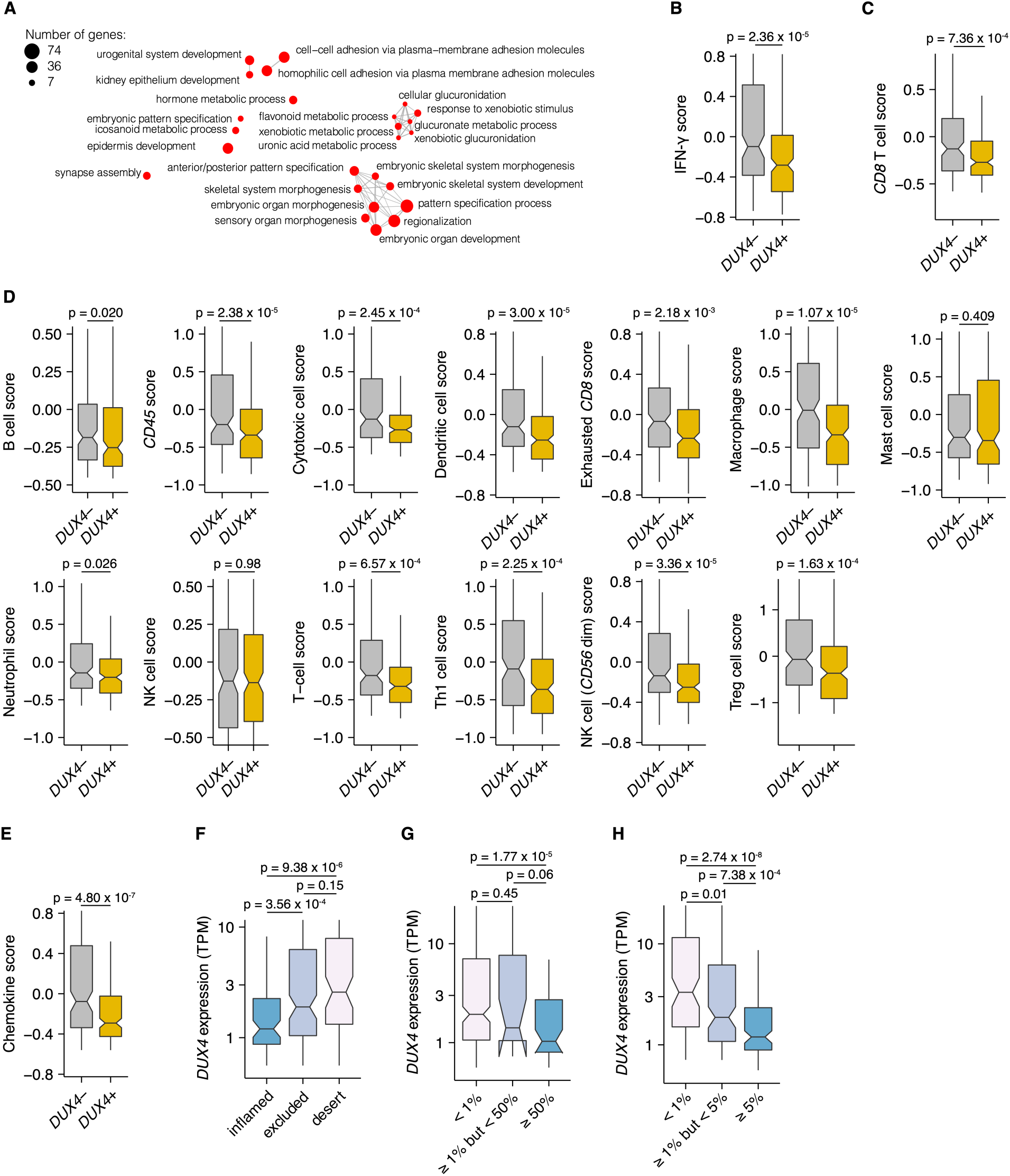
*DUX4*-positivity is correlated with an embryonic gene expression signature, downregulation of interferon-gamma signaling, and exclusion of diverse immune cell types. (**A**) As in (Figure 2A), but the Gene Ontology (GO) enrichment network analysis corresponding to *DUX4*-upregulated genes, compared against the set of coding genes, is shown. (**B**) Interferon-gamma (IFN-γ) signature score (Ayers et al., 2017). The *p*-value was estimated via a Mann-Whitney *U* test. (**C**) CD8 T cell score from (Danaher et al., 2017). The *p*-value was estimated via a Mann-Whitney *U* test. (**D**) As in (**C**), showing the other immune cell signatures available in (Danaher et al., 2017). (**E**) Chemokine signature score (Coppola et al., 2011). The *p*-value was estimated via a Mann-Whitney *U* test. (**F**) *DUX4* expression (TPM) in inflamed, immune excluded, and immune desert tumors. The phenotypes are based on CD8^+^ T cell abundance and degree of tumor infiltration determined by anti-CD8 staining of tumor FFPE sections in the original study (Mariathasan et al., 2018). The *p*-values were estimated via the Mann-Whitney *U* test. (**G**) *DUX4* expression (TPM) in advanced urothelial carcinoma tumors. The percentage of tumor cells with positive PD-L1 staining are indicated on the x-axis. The *p*-values were estimated via the Mann-Whitney *U* test. (**H**) As in (**G**), but showing the percentage of tumor-infiltrating immune cells (lymphocytes, macrophages, and dendritic cells) with positive PD-L1 staining on the x-axis.

**Figure S3.**
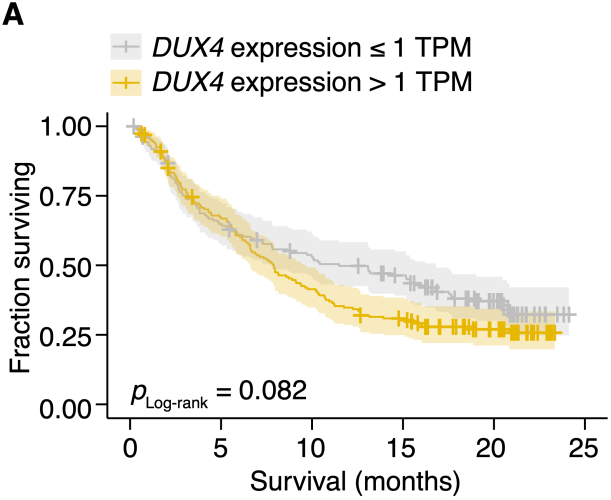
*DUX4* expression status stratifies patients according to survival. (**A**) Kaplan-Meier (KM) estimates of overall survival (solid lines), 95% confidence intervals (transparent ribbons), and censored events (crosses) for ICI-treated advanced urothelial carcinoma patients stratified by *DUX4* expression status. The *p*-value was estimated via a log-rank test.

**Figure S4.**
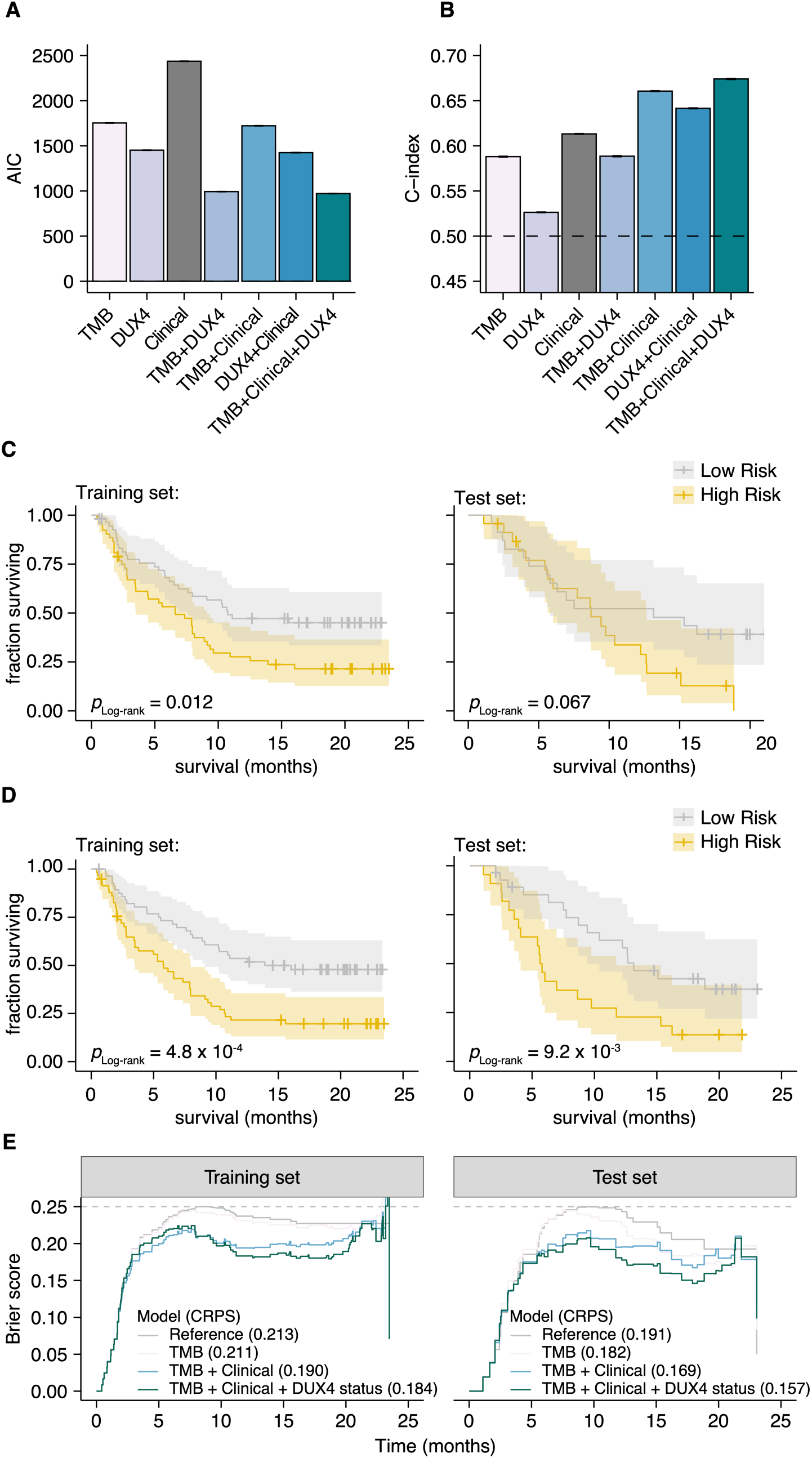
Cox Proportional Hazards regression models containing *DUX4* expression status as a predictor have a better fit to the data. (**A**) Akaike information criterion (AIC) measurements for goodness of fit for the full (TMB, Clinical, DUX4 expression) vs. reduced Cox PH models, where lower values indicate better fit. The bootstrapped AIC mean and the 95% confidence interval are illustrated. Clinical (ECOG Performance Status and Platinum treatment history). (**B**) Harrell’s concordance indices (C-index) for the full (TMB, Clinical, DUX4 expression) vs. reduced Cox PH models, where high values indicate better model performance. The bootstrapped C-index mean and the 95% confidence interval are illustrated. (**C**) Kaplan-Meier (KM) estimates of overall survival, 95% confidence interval (transparent ribbon), and censored events (crosses) for low-risk (solid gray line) and high-risk (solid orange line) patients in the training (left) and test (right) sets. Risk group assignments were based on risk scores estimated by the Cox PH model with only TMB as a predictor. *p*-values were estimated via a log-rank test. (**D**) As in (C), but the risk group assignments were based on risk scores estimated by the Cox PH model with TMB, ECOG Performance Status, and Platinum treatment history as predictors. (**E**) Time-dependent Brier scores for the full and reduced Cox PH models applied on the training (left) and test (right) sets. The Continuous Ranked Probability Scores (CRPS), defined as the integrated Brier score divided by time, are shown in parentheses. Reference refers to the Kaplan-Meier prediction model. A Brier score = 0.25 indicates random guessing (gray dashed line).

**Figure S5.**
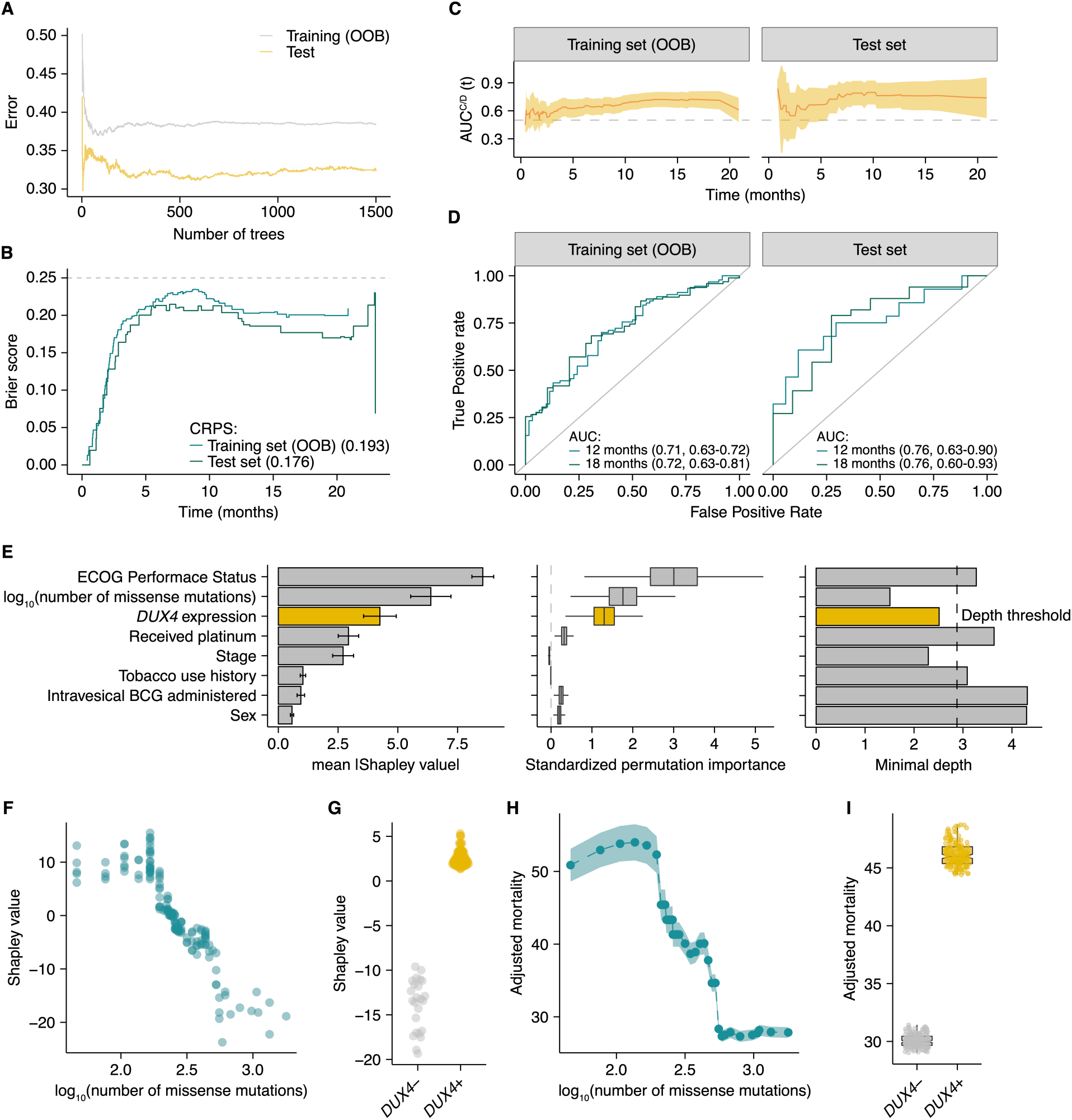
A Random Survival Forest model quantifies the effect of *DUX4* status on overall survival probability in the context of immune checkpoint inhibition. (**A**) Error (1 – Harrell’s concordance index) as a function of the number of trees in the Random Survival Forest (RSF) model. The training out-of-bag error (OOB error, solid gray line) and the test error (solid orange line) are shown. 1500 trees were used in the final model. (**B**) Time-dependent Brier scores for the RSF model estimated from the training (solid turquoise line) or test (solid teal line) sets. OOB survival predictions were used to calculate the Brier score for the training set. The Continuous Ranked Probability Scores (CRPS), defined as the integrated Brier score divided by time, for both sets are shown. A Brier score = 0.25 indicates random guessing (gray dashed line). (**C**) Time-dependent ROC analyses. The AUC^C/D^ (solid orange line) and the 95% confidence interval (transparent orange ribbon) are shown over the observation time for the training and test sets. The OOB mortality predictions were used to calculate the training set AUC^C/D^. (**D**) The Receiver Operating Characteristic (ROC) curves for the RSF model at 12 (solid turquoise line) and 18 (solid teal line) months. The Cumulative/Dynamic Area Under the ROC Curve (AUC^C/D^) and 95% confidence interval are specified. The OOB mortality predictions were used to calculate the training set AUC^C/D^. (**E**) RSF feature importance. The mean absolute Shapley values and the 95% confidence intervals are shown (left). The standardized permutation importance and confidence regions estimated via delete-*d* jackknife subsampling are plotted (middle); noise variables are indicated by permutation importance measures ≤ 0. The minimal depth measures are shown (right); variables with values exceeding the depth threshold (gray dashed line) are designated as noise variables. (**F**) Shapley dependence plot illustrating the relationship between tumor mutational burden (TMB, number of missense mutations) and mortality. Each point corresponds to a single patient. (**G**) As in (**F**), but showing *DUX4* expression status. (**H**) Partial plot illustrating the marginal effect of TMB on mortality. Each point corresponds to the average RSF mortality predictions when TMB is fixed to the indicated value for all patients. The transparent ribbon corresponds to the 95% confidence interval.(I) Partial plot illustrating the marginal effect of *DUX4* expression status. The points correspond to the RSF prediction for mortality for each patient when *DUX4* expression status is fixed to the indicated value for the entire cohort.

## SUPPLEMENTAL FILES

**Table S1.** Cox Proportional Hazards Regression for Overall Survival (TGFB1 expression included)

| Characteristic | Univariate |  |  |  | Multivariate<sup>a,b</sup> |  |  |  |
| --- | --- | --- | --- | --- | --- | --- | --- | --- |
|  | HR | 95% CI | p-value | q-value<sup>c</sup> | HR | 95% CI | p-value | q-value<sup>c</sup> |
| Tumor mutational burden<sup>d</sup> | 0.43 | 0.28 - 0.66 | $1.10 \times 10^{-4}$ | $8.27 \times 10^{-4}$ | 0.14 | 0.06 - 0.30 | $8.41 \times 10^{-7}$ | $1.26 \times 10^{-5}$ |
| DUX4 expression: |  |  |  |  |  |  |  |  |
| < 0.25 TPM (reference) | — | — | — | — | — | — | — | — |
| > 1 TPM | 1.85 | 1.10 - 3.11 | 0.021 | 0.079 | 3.12 | 1.52 - 6.38 | $1.87 \times 10^{-3}$ | $7.01 \times 10^{-3}$ |
| ECOG Performance Status<sup>e</sup>: |  |  |  |  |  |  |  |  |
| 0 (reference) | — | — | — | — | — | — | — | — |
| 1 | 2.10 | 1.58 - 2.79 | $2.95 \times 10^{-7}$ | $4.43 \times 10^{-6}$ | 2.86 | 1.79 - 4.57 | $1.18 \times 10^{-5}$ | $8.84 \times 10^{-5}$ |
| 2 | 1.97 | 1.04 - 3.73 | 0.036 | 0.091 | 5.31 | 1.79 - 15.7 | $2.58 \times 10^{-3}$ | $7.73 \times 10^{-3}$ |
| Received platinum: |  |  |  |  |  |  |  |  |
| No (reference) | — | — | — | — | — | — | — | — |
| Yes | 1.41 | 1.01 - 1.98 | 0.047 | 0.10 | 2.69 | 1.47 - 4.93 | $1.32 \times 10^{-3}$ | $6.61 \times 10^{-3}$ |
| Intravesical BCG administered: |  |  |  |  |  |  |  |  |
| No (reference) | — | — | — | — | — | — | — | — |
| Yes | 0.98 | 0.73 - 1.32 | 0.89 | 0.92 | 0.64 | 0.38 - 1.06 | 0.08 | 0.15 |
| Tobacco use history: |  |  |  |  |  |  |  |  |
| Never (reference) | — | — | — | — | — | — | — | — |
| Previous | 0.90 | 0.68 - 1.19 | 0.45 | 0.61 | 1.21 | 0.78 - 1.87 | 0.39 | 0.45 |
| Current | 1.02 | 0.65 - 1.61 | 0.92 | 0.92 | 1.43 | 0.76 - 2.67 | 0.27 | 0.36 |
| Stage: |  |  |  |  |  |  |  |  |
| I | — | — | — | — | — | — | — | — |
| II | 0.96 | 0.69 - 1.35 | 0.83 | 0.92 | 1.61 | 0.91 - 2.86 | 0.10 | 0.17 |
| III | 1.23 | 0.86 - 1.75 | 0.25 | 0.38 | 1.62 | 0.96 - 2.74 | 0.07 | 0.15 |
| IV | 0.98 | 0.68 - 1.42 | 0.92 | 0.92 | 0.46 | 0.24 - 0.88 | 0.020 | 0.050 |
| Sex: |  |  |  |  |  |  |  |  |
| Male (reference) | — | — | — | — | — | — | — | — |
| Female | 1.23 | 0.91 - 1.66 | 0.18 | 0.31 | 0.76 | 0.47 - 1.23 | 0.26 | 0.36 |
| TGFB1 expression: |  |  |  |  |  |  |  |  |
| ≤ 25<sup>th</sup> percentile (Q1, reference) | — | — | — | — | — | — | — | — |
| > 25<sup>th</sup> percentile and ≤ 50<sup>th</sup> percentile (Q2) | 1.30 | 0.88 - 1.91 | 0.19 | 0.31 | 0.75 | 0.40 - 1.41 | 0.38 | 0.45 |
| > 50<sup>th</sup> percentile and ≤ 75<sup>th</sup> percentile (Q3) | 1.52 | 1.04 - 2.23 | 0.031 | 0.091 | 1.04 | 0.55 - 1.95 | 0.90 | 0.91 |
| > 75<sup>th</sup> percentile (Q4) | 1.68 | 1.16 - 2.45 | $6.46 \times 10^{-3}$ | 0.032 | 0.96 | 0.50 - 1.87 | 0.91 | 0.91 |

<sup>a</sup>Log-rank test: p-value = $1.27 \times 10^{-7}$ <sup>b</sup>Akaike Information Criterion = 975.30; Bayesian Information Criterion = 1015.67; Harrell's Concordance Index = 0.71<sup>c</sup>Benjamini-Hochberg FDR correction<sup>d</sup>log<sub>10</sub>(number of missense mutations)<sup>e</sup>Eastern Oncology Cooperative Group Performance Status

